# Rare longevity-associated variants, including a reduced-function mutation in cGAS, identified in multigenerational long-lived families

**DOI:** 10.64898/2025.12.04.689698

**Authors:** Pasquale C. Putter, Di Guan, Thies Gehrmann, Daniel Kolbe, Jiping Yang, HyeRim Han, Seungsoo Kim, Nico Lakenberg, H. Eka D. Suchiman, Stella Trompet, Georg C. Liu, Eugen Ballhysa, Adam Antebi, Niels M.A. van den Berg, Almut Nebel, Marian Beekman, Yousin Suh, P. Eline Slagboom, Joris Deelen

## Abstract

Life expectancy has steadily increased in the last two centuries, while healthspan has been lagging behind. Survival into extreme ages strongly clusters within families which often exhibit a delayed onset of (multi)morbidity, yet the underlying protective genetic mechanisms are still largely undefined. We performed affected sib-pair linkage analysis in 212 sibships enriched for ancestral longevity and identified four genomic regions (LOD_max_ ≥3.0) at *1q21*.*1*, *6p24.3, 6q14.3,* and *19p13.3*. Within these regions, we prioritized 12 rare protein-altering variants in seven candidate genes (*NUP210L, SLC27A3, CD1A, CGAS, IBTK, RARS2,* and *SH2D3A*) located in longevity-associated loci. Notably, a missense variant in *CGAS* (rs200818241), was present in two sibships. Using human- and mouse-based cell models, we showed that rs200818241 reduced protein stability and attenuated activation of the canonical cGAS-STING pathway in a cell-type specific manner. This dampened signalling mitigated inflammation and delayed cellular senescence, mechanisms that may contribute to the survival advantage of *CGAS* variant carriers. Our findings indicate novel rare variants and candidate genes linked to familial longevity and highlight the cGAS-STING pathway as a potential contributor to the protective mechanisms underlying human longevity.

## Main

In the last 200 years, human life expectancy has doubled in developed nations, but healthspan (i.e. the number of disease-free years lived) has not kept pace with lifespan [1]. Chronic disease incidence rises with age, leading to multi-morbidity and an increasing social and economic burden [2, 3]. Lifespan is influenced by many factors such as education, socioeconomic status, lifestyle, and, to some extent, genetics. Early studies estimated the heritability of lifespan at 10–25% [4, 5], but these studies did not distinguish between deaths from intrinsic biological processes and deaths from extrinsic causes such as accidents or infections. Recent modelling correcting for extrinsic mortality and age cutoffs suggests that the heritability of lifespan due to intrinsic processes may likely be above 50% [6].

The genetic influence is assumed to be even stronger for longevity, defined as survival to an extreme old age, as it clusters within families [7]. Familial longevity is transmitted as a quantitative trait for families in which members who survive into the top 10% of their birth cohort. Every ancestor meeting this criterium contributes to the additional survival probability of their descendants [8]. Members of long-lived families, such as those included in the Leiden Longevity Study (LLS) [9], exhibit a lifelong survival advantage, and a delayed onset of age-related multimorbidity. At the biological level, they display favourable molecular health profiles in mid-life, and their fibroblasts demonstrate an enhanced stress response capacity [10–13].

Genetic studies of longevity aim to uncover genes and pathways underlying these systemic benefits. So far, most work has focused on common genetic variants in unrelated individuals. Only a limited number of robust associations have been identified, such as *APOE* and *FOXO3* [14, 15], which are insufficient to fully explain the remarkable health and survival advantage of long-lived families. Hence, the missing heritability of (familial) longevity may lie in rare protective genetic variants with larger effects [16–19]. Indeed, functional analysis of rare variants identified by candidate-gene approaches have shown impact on conserved pathways of longevity in cellular models [20, 21]. Yet, such approaches are limited, as they may miss rare variants in genes without a priori evidence for a role in longevity.

Here, we applied a hypothesis-free genetic linkage analysis in LLS families enriched for ancestral longevity [22]. We identified novel linkage regions and prioritized high-impact, protein-altering rare genetic variants associated with longevity, including a missense variant in *CGAS*, which we functionally characterized to reveal its effects on protein stability, cellular senescence, and on the canonical cGAS-STING signalling pathway.

## Results

### Sibships enriched for ancestral longevity score better on lifespan, well-being, and cognition than community-dwelling controls

To identify rare genetic variants contributing to familial longevity, we performed linkage analysis using long-lived sibships from the LLS cohort consisting of Dutch families with a history of longevity across multiple generations. The LLS was initiated with 421 nonagenarian sibships enrolled between 2002-2006, where men and women had to be older than 89 and 91 years respectively, thereby representing 0.5% of the Dutch population at that time. To make the familial longevity phenotype more restrictive to heritable influences, we used the recently created ancestral definition of familial longevity based on at least a parent and at minimum two nonagenarian siblings belonging to the top decile of their respective sex-specific birth cohort [22]. This resulted in a selection of 212 (50.4%, N=479) out of 421 sibships for the genetic linkage analysis. These selected LLS sibships reached a mean lifespan of 97.4 years (SD ± 3.5) (**Table 1**).

**Table 1.** Descriptive statistics of the selected Leiden Longevity Study (LLS) sibships enriched for ancestral longevity.

| Trait | Characteristics |
| --- | --- |
| Long-lived siblings, N | 479 |
| Sibships, N | 212 |
| Sibship size, mean (range) | 2.4 (2-4) |
| Females, N (%) | 295 (61.6) |
| Lifespan in years, mean (range) (SD) | 97.6 (89.7 – 109) (3.5) |

To investigate how the selected longevity-enriched sibships from the LLS perform on common geriatric assessments we compared them to a subset of community-dwelling Dutch individuals from the Leiden 85+ study that reached an age of at least 90 years (N=277) [23]. These individuals are culturally and genetically very similar to the selected LLS sibships and, moreover, they were born around the same time, making them a strong control group. The selected LLS sibships showed a significantly longer lifespan and scored better on cognitive function and subjective well-being than the nonagenarian subset of individuals from the Leiden 85+ study, while no differences were observed for (instrumental) activities of daily living, which refer to the ability to perform everyday personal care and independent functioning (**Extended data 1, Supplementary Table 1**). Of note, before dichotomizing the (instrumental) activities of daily living scores, the selected LLS sibships appeared to have a distribution shifted more toward the healthier end, indicating they are better capable of living independently compared to the nonagenarian individuals from the Leiden 85+ study (**Extended data 2**).

### Sib-pair linkage analysis identifies four novel genetic regions contributing to familial longevity

We subsequently performed a non-parametric genome-wide affected sib-pair (ASP) linkage analysis on the 212 selected long-lived LLS sibships. Significant evidence for linkage (logarithm of the odds (LOD)-score≥3) was observed at four chromosomal regions (**Figure 1A**), defined by the peak maximum (LOD_max_) and the surrounded LOD-score>2 region, namely *Chr1q21.1*, *Chr6p24.3*, *Chr6q14.3*, and *Chr19p13.3* (**Figure 1 B-D**, **Table 2**), spanning 35.7 Mb and harbouring 373 genes (**Supplementary Table 2**). Three of these regions (*Chr1q21.1*, *Chr6p24.3*, and *Chr6q14.4*) have not yet been linked to familial longevity, while the one on chromosome 19 (*Chr19p13.3*) has a small region that partially overlaps with a locus found in an earlier study using long-lived sibships from families across Europe [17].

**Figure 1.**
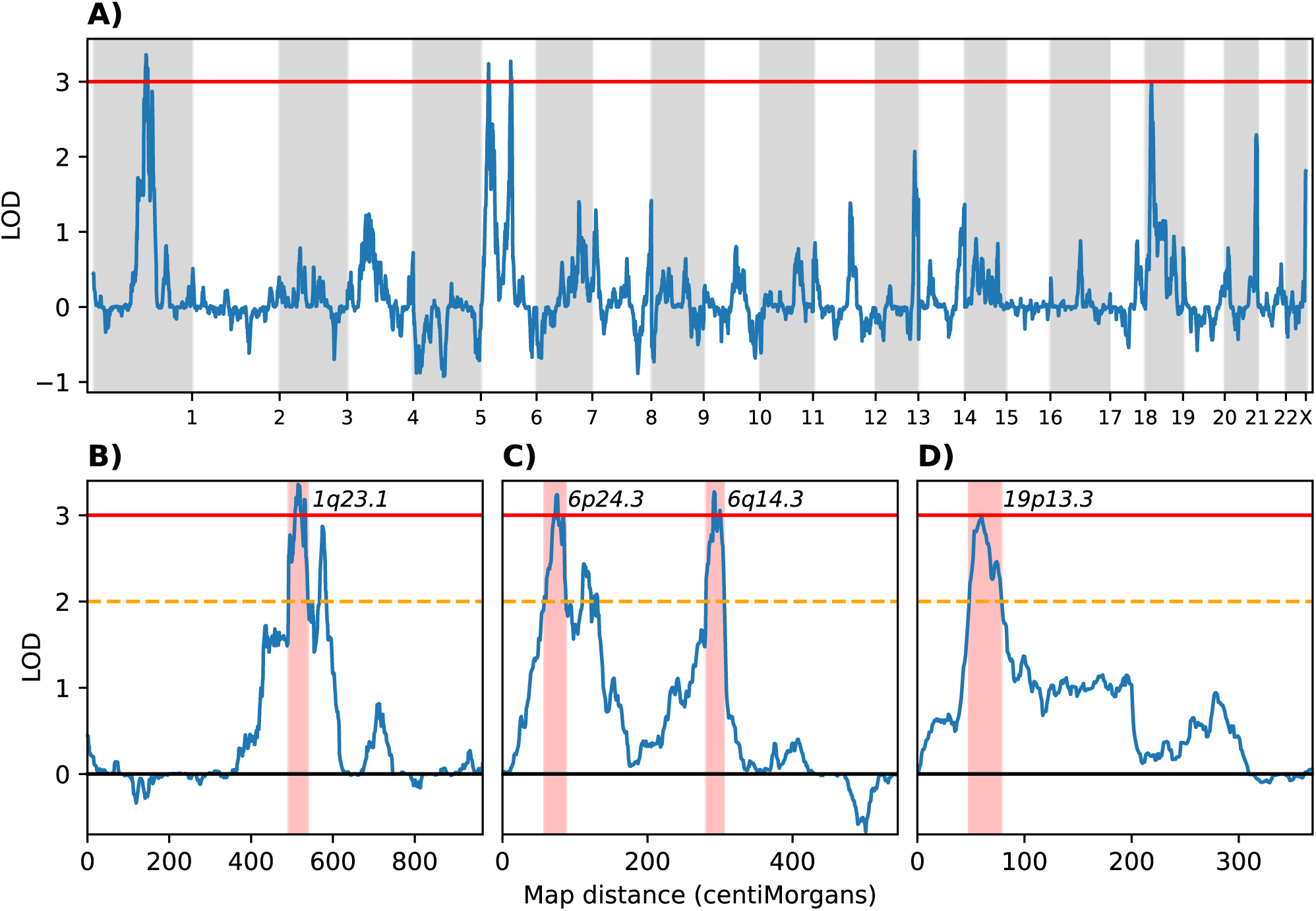
Genome-wide linkage analysis identifies four loci linked to ancestral familal longevity. **A)** Genome-wide non-parametric linkage (NPL) analysis in 212 selected LLS sibships enriched for ancestral longevity showing logarithm of the odds (base 10) (LOD)-scores across the genome. The X-axis shows the chromosomal position in centiMorgan (cM) and the Y-axis shows the LOD-score. The solid blue line displays the NPL LOD-scores. The solid red line indicates the genome-wide significance threshold (LOD-score=3), and the dashed orange line represents the suggestive threshold (LOD-score=2). **B-D)** Regional linkage peaks showing evidence for linkage on **B)** Chromosome 1, **C)** Chromosome 6, **D)** Chromosome 19. Red highlighted area indicate the regions of primary interest under the peaks. Cytoband labels mark the locations of the maximum LOD-scores (LODmax), which are used to name the linked loci.

**Table 2.** Description of four chromosomal regions linked to familial longevity.

| Peak | $LOD_{max}$ | Region of interest<br>($LOD\text{-score} > 2$) | Interval | Locus | SNP | Location |
| --- | --- | --- | --- | --- | --- | --- |
| **Chr1q23.1** | 3.3 at<br>rs1864348 | 492-538cM | Start | chr1q21.3 | rs4845364 | 153,294,652 |
|  |  |  | End | chr1q23.3 | rs1832739 | 162,210,470 |
| **Chr6p24.3** | 3.2 at<br>rs384323 | 58-87cM | Start | chr6p24.2 | rs3018 | 5,723,117 |
|  |  |  | End | chr6p25.1 | rs4713064 | 11,198,497 |
| **Chr6q14.3** | 3.3 at<br>rs4574603 | 281-305cM | Start | chr6q13 | rs1767861 | 73,578,414 |
|  |  |  | End | chr6q15 | rs4707657 | 92,254,619 |
| **Chr19p13.3** | 3.0 at<br>rs673153 | 49-78cM | Start | chr19p13.2 | rs2779180 | 4,610,237 |
|  |  |  | End | chr19p13.3 | rs2042902 | 7,278,451 |

**LOD:** Logarithm of the odds (base 10); **$LOD_{max}$** highest LOD-score reaching significance ( $LOD \geq 3$ ) in region of interest; **Location:** Location in base pairs according to Grch37 assembly.

### Collapsing filtering strategy leads to identification of novel rare functional variants linked to familial longevity

#### Common variants

Within these four identified linkage regions there were no common genome-wide significant single-nucleotide polymorphisms (SNPs; P-value ≤ 5·10^-8^) associated with longevity or parental lifespan as reported in the GWAS Catalog (v1.0.3.1 – 2025-06-16) [24], although we identified 10 suggestive significant SNPs (P-value ≤ 1·10^-6^) previously associated with these traits (**Supplementary Table 3**). Moreover, we performed a genetic association analysis within the LLS, using imputed SNP microarray data, comparing the selected LLS long-lived sibships to controls from the general population, i.e. the middle-aged partners of their offspring with no history of familial longevity. Within the predefined linkage regions of interest, no SNPs reached the suggestive significance threshold (P-value ≤ 1·10^-5^; **Extended Data 3, Supplementary Table 4**). Additionally, we did not identify any associative signal of the previously reported GWAS Catalog SNPs within the LLS (**Supplementary Table 3**). Looking across the entire genome, the top association signal was observed at rs754579 (P-value = 2.43·10^-7^), located within the intronic regions of a lncRNA near *TUB3AC*. Moreover, we replicated the well-known longevity association of rs429358 in the *APOE* locus (P-value = 5.20·10^-7^) [15]. Although these results did not reach genome-wide significance, which is likely attributable to the small sample size of our cohort, it shows that we should have been able to identify common genetic variants located under the linkage peaks with substantial effects on longevity. Altogether this supports the idea that (familial) longevity in the LLS may primarily be driven by rare genetic variants.

#### Rare variants

To identify rare variants within the four linkage regions of interest, we obtained whole-genome sequencing data for one sibling from each sibship in 179 of the 212 long-lived families [25]. Since we lacked a large enough control group of non-long-lived individuals with sequencing data using the same platform, we were unable to perform a burden analysis to detect enrichment of rare variants in specific genes located under the linkage peaks. Instead, we performed a collapsing filtering variant strategy (**Extended Data 4**). We focused on protein-altering variants with a high predicted impact (Combined Annotation Dependent Depletion (CADD) score >15), located in loci that were identical-by-descent (IBD)≥1 between siblings, and that were absent in our Dutch control cohort (N=100) and very rare (Minor Allele Frequency; MAF<0.2%) in the general non-Finnish European population available in the Genome Aggregation Database (gnomAD) v2.1.1 [26]. We subsequently continued with variants that were shared between multiple long-lived families or that were located in a gene in which we identified multiple variants in different long-lived families. These variants, all of which were present in the heterozygous state in the identified carriers, were subsequently validated using Sanger sequencing in the identified carrier and at least one sibling (**Supplementary Table 6**). Next, we checked the frequency of the selected variants in an independent cohort of German long-lived individuals (LLI) and younger controls, who are of a similar ethnic background but are not known to be enriched for ancestral longevity. We excluded variants that were enriched in the controls in comparison to the LLI, since these are less likely to contribute to (familial) longevity. This strategy yielded 12 variants residing in seven genes (**Table 3**). Two of the identified variants were present in two independent long-lived families, i.e. rs200818241 (**Extended data 5A)** in *CGAS*, and rs146711285 in *SH2D3A*.

**Table 3.**
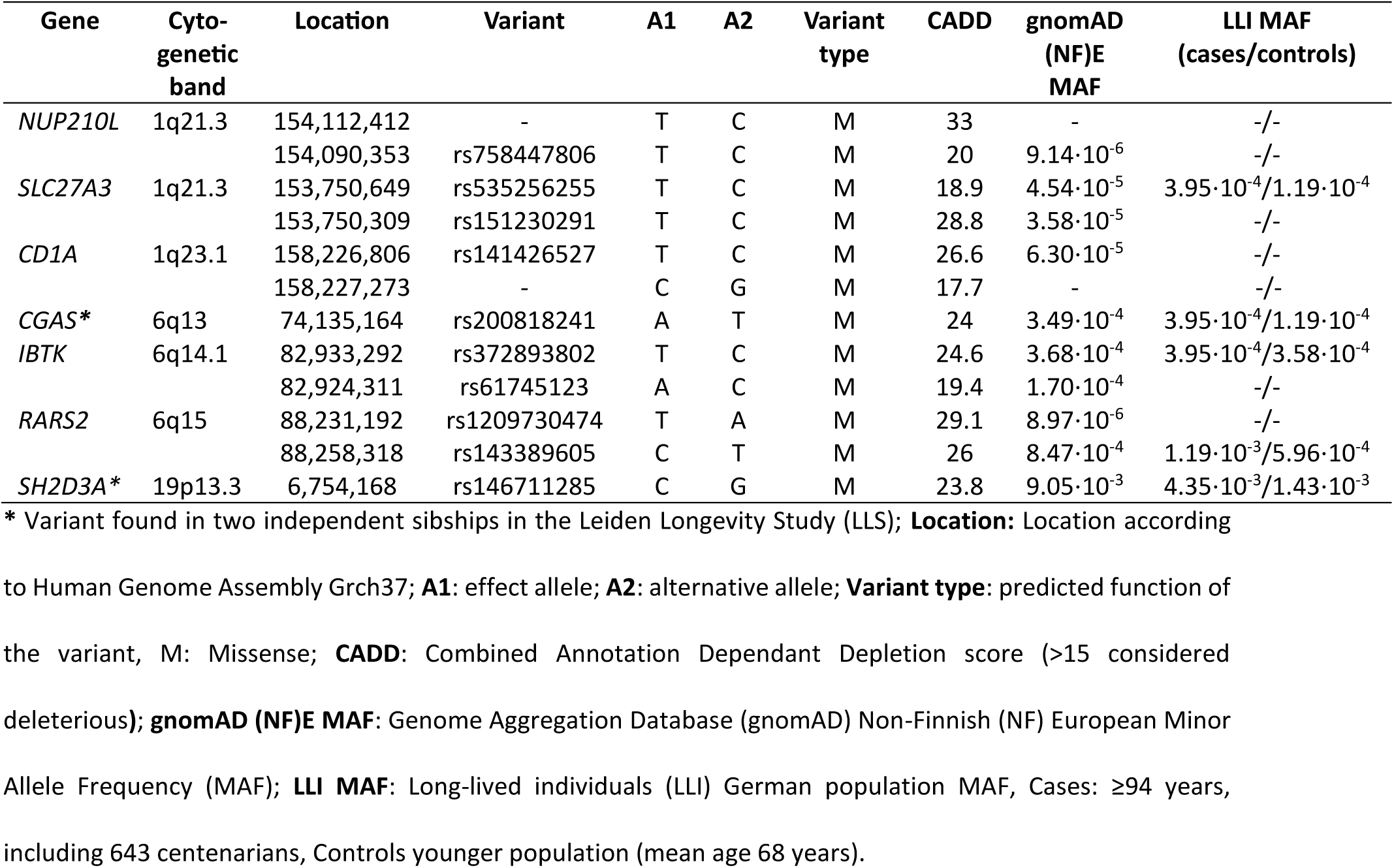
Positional rare candidate variants potentially contributing to familial longevity.

| Gene | Cyto-genetic band | Location | Variant | A1 | A2 | Variant type | CADD | gnomAD (NF)E MAF | LLI MAF (cases/controls) |
| --- | --- | --- | --- | --- | --- | --- | --- | --- | --- |
| *NUP210L* | 1q21.3 | 154,112,412 | - | T | C | M | 33 | - | -/- |
| | | 154,090,353 | rs758447806 | T | C | M | 20 | $9.14 \cdot 10^{-6}$ | -/- |
| *SLC27A3* | 1q21.3 | 153,750,649 | rs535256255 | T | C | M | 18.9 | $4.54 \cdot 10^{-5}$ | $3.95 \cdot 10^{-4} / 1.19 \cdot 10^{-4}$ |
| | | 153,750,309 | rs151230291 | T | C | M | 28.8 | $3.58 \cdot 10^{-5}$ | -/- |
| *CD1A* | 1q23.1 | 158,226,806 | rs141426527 | T | C | M | 26.6 | $6.30 \cdot 10^{-5}$ | -/- |
|  |  | 158,227,273 | - | C | G | M | 17.7 | - | -/- |
| *CGAS\** | 6q13 | 74,135,164 | rs200818241 | A | T | M | 24 | $3.49 \cdot 10^{-4}$ | $3.95 \cdot 10^{-4} / 1.19 \cdot 10^{-4}$ |
| *IBTK* | 6q14.1 | 82,933,292 | rs372893802 | T | C | M | 24.6 | $3.68 \cdot 10^{-4}$ | $3.95 \cdot 10^{-4} / 3.58 \cdot 10^{-4}$ |
| | | 82,924,311 | rs61745123 | A | C | M | 19.4 | $1.70 \cdot 10^{-4}$ | -/- |
| *RARS2* | 6q15 | 88,231,192 | rs1209730474 | T | A | M | 29.1 | $8.97 \cdot 10^{-6}$ | -/- |
| | | 88,258,318 | rs143389605 | C | T | M | 26 | $8.47 \cdot 10^{-4}$ | $1.19 \cdot 10^{-3} / 5.96 \cdot 10^{-4}$ |
| *SH2D3A\** | 19p13.3 | 6,754,168 | rs146711285 | C | G | M | 23.8 | $9.05 \cdot 10^{-3}$ | $4.35 \cdot 10^{-3} / 1.43 \cdot 10^{-3}$ |

\* Variant found in two independent sibships in the Leiden Longevity Study (LLS); **Location:** Location according to Human Genome Assembly Grch37; **A1:** effect allele; **A2:** alternative allele; **Variant type:** predicted function of the variant, M: Missense; **CADD:** Combined Annotation Dependant Depletion score (>15 considered deleterious); **gnomAD (NF)E MAF:** Genome Aggregation Database (gnomAD) Non-Finnish (NF) European Minor Allele Frequency (MAF); **LLI MAF:** Long-lived individuals (LLI) German population MAF, Cases: ≥94 years, including 643 centenarians, Controls younger population (mean age 68 years).

### *CGAS* expression is altered in different tissues with age

We focused on rs200818241 because *CGAS* has been implicated in ageing through inflammation, neurodegeneration and promotion of the senescent-associated secretory phenotype (SASP) in mouse models [27–29]. Blocking the cGAS-STING pathway has also been proposed as a strategy to slow age-related degenerative processes [30]. Using data from the Genotype-Tissue Expression (GTEx) consortium [31], we found that *CGAS* expression showed tissue-specific age effects, decreasing in blood and visceral adipose but increasing in adrenal, breast, esophagus, lung, nerve prostate, and skin (**Supplementary Table 7**), supporting a potential role for *CGAS* in human ageing. Different cohorts have reported in whole blood both up- and downregulation with age, suggesting that the role of cGAS is context- and age-dependent and warrants further research [32]. To test whether rs200818241 directly affect *CGAS* expression, we performed allele-specific expression analysis in blood of both the long-lived individuals themselves and their variant-carrying middle-aged offspring. No allele-specific expression was observed (**Extended data 5B**), indicating that rs200818241 does not alter *cis-*regulatory activity in blood.

### rs200818241 suppresses cGAS-STING activation and reduces cGAS protein stability

The rs200818241 variant introduces a thymine-to-adenine base change, resulting in an aspartate-to-valine substitution at position 452 (D452V) of the cGAS protein (**Extended data 5C**). This substitution occurs towards the C-terminal end in the Mab21 domain, a region critical for cGAS catalytic activity and nucleic acid binding of the protein [33]. To assess its functional impact, we expressed the mutant (cGAS-rs200818241) or wild-type (cGAS-WT) forms of cGAS in different human cell lines using viral vectors. Stress-induced cGAS activation normally increases phosphorylation of STING, TBK1, and IRF3, inducing type I interferons (e.g., *IFNB1*), interferon-stimulated genes (e.g., *IFI44*), and pro-inflammatory cytokines (e.g., *IL6*, *CXCL8*, *CCL2*, *CXCL1*, and *CXCL10*) [34].

In human mesenchymal stromal cells (hMSCs), the cGAS-rs200818241 variant resulted in significantly lower expression of downstream target genes compared to cGAS-WT following viral transduction (**Figure 2A**). Consistently, phosphorylation of TBK1 and IRF3 were markedly reduced in cells expressing cGAS-rs200818241 relative to cGAS-WT (**Figure 2B**). These findings indicate that rs200818241 variant leads to attenuated cGAS-STING signalling pathway activation in response to cytosolic DNA. Similar effects were observed in granulosa cells (KGN) (**Figure 2C-D**) and primary human astrocytes (**Extended data 6A**). However, in fibroblasts, the effect was partial and only *IFI44*, and *CXCL8*, showed a similar pattern as the other cell types (**Extended data 6B**). Together, these findings demonstrate that rs200818241 attenuates the activation of the canonical cGAS-STING pathway in a cell type-specific manner.

**Figure 2.**
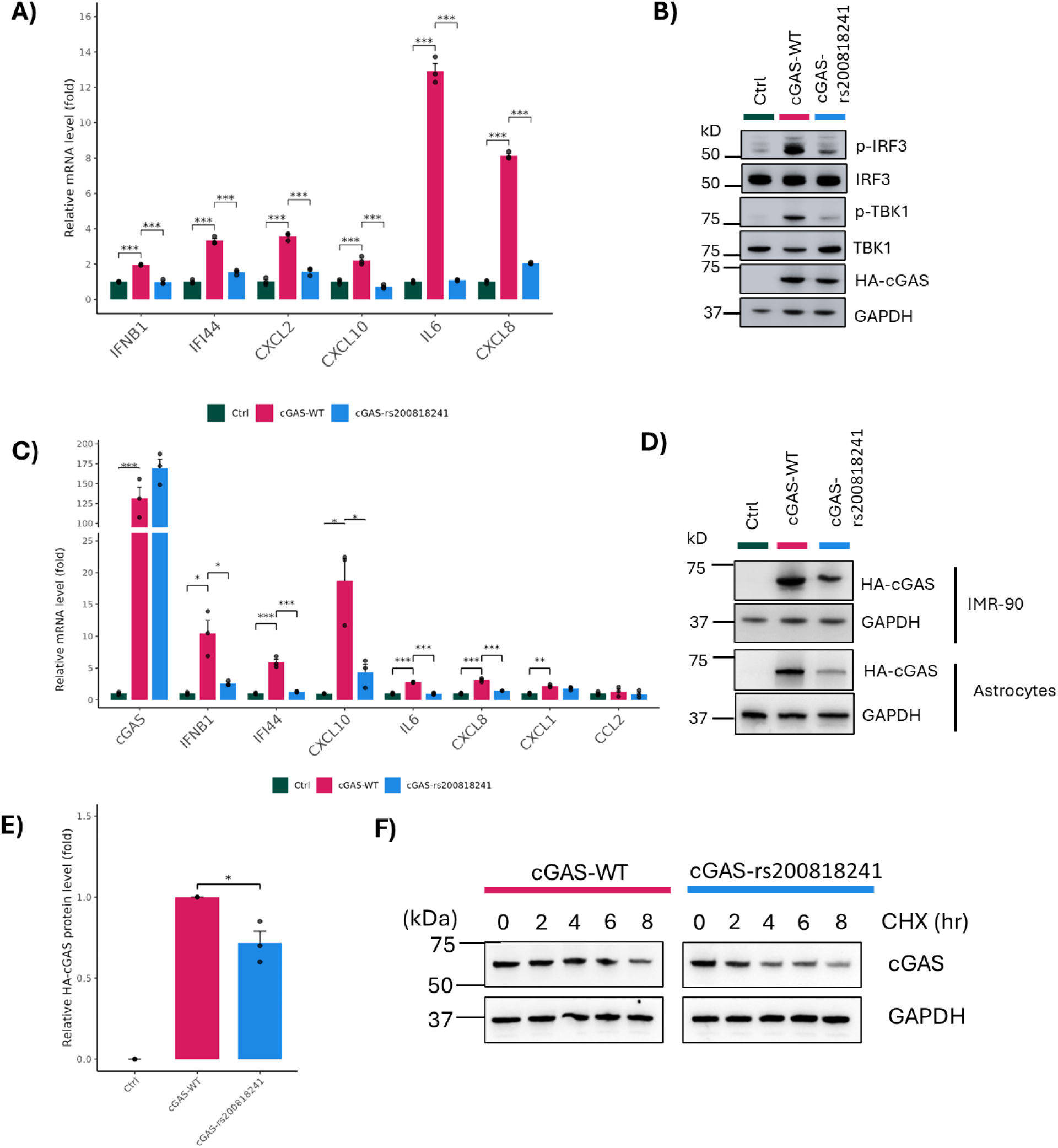
Longevity-associated rare coding variant rs200818241 in *CGAS* suppresses activation of the canonical cGAS-STING pathway and reduces protein stability in human cell lines. **A-B**) Quantitative PCR (qPCR) analysis of downstream gene expression (**A**) and western blot analysis of phosphorylated mediators (**B**) in the canonical cGAS-STING pathway in human mesenchymal stromal cells (hMSCs) expressing the longevity-associated cGAS variant (cGAS- rs200818241), wild-type cGAS (cGAS-WT), or an empty vector control (Ctrl). **C**), qPCR analysis of downstream gene expression in the canonical cGAS-STING pathway in human granulosa (KGN) cells expressing cGAS-rs200818241, cGAS-WT, or Ctrl. **D**), Western blot analysis of cGAS protein levels in fibroblasts (IMR-90) and primary human astrocytes expressing cGAS-rs200818241, cGAS-WT, or Ctrl. **E**), Quantification of cGAS protein levels in hMSCs, normalized to GAPDH, showing reduced levels in cells expressing cGAS-rs200818241 compared to cGAS-WT. **F**), Cycloheximide (CHX) chase assay showing faster degradation of the cGAS protein in hMSCs expressing cGAS-rs200818241 compared to cGAS-WT. All experiments were performed in triplicate. Statistical significance is denoted as follows: *p < 0.05, **p < 0.01, ***p < 0.001.

To investigate the mechanism by which rs200818241 suppresses activation of the canonical cGAS-STING pathway, we compared the expression levels of cGAS in cells expressing cGAS-rs200818241 versus cGAS-WT. While *CGAS* mRNA levels were comparable between the two (**Figure 2C**), the protein levels of cGAS were consistently reduced in hMSCs, IMR-90, and primary human astrocytes expressing cGAS-rs200818241 (**Figure 2B,D-E**), suggesting that rs200818241 decreases cGAS protein stability. To directly assess the impact of rs200818241 on protein stability, we expressed both cGAS-rs200818241 and cGAS-WT in hMSCs and performed a cycloheximide (CHX) chase assay to evaluate protein degradation rates. Following CHX treatment, cGAS-rs200818241 degraded more rapidly than cGAS-WT (**Figure 2F**), providing further evidence that rs200818241 reduces protein stability, which likely underlies its attenuated activation of the canonical cGAS-STING pathway.

### rs2008182410 in cGAS extends cellular lifespan

Given that reduced activation of the canonical cGAS-STING pathway has been associated with mitigation of ageing-related processes in mice, including protection against cognitive decline [27, 35], we tested whether rs200818241 influences replicative lifespan and senescence in hMSCs. We observed that cells expressing the cGAS-WT rapidly enter a state of senescence, characterized by proliferation arrest (**Figure 3A**), activation of the canonical cGAS-STING pathway and increased p16 expression (**Figure 3B**), and accumulation of senescence-associated β-galactosidase (SA-β-gal) positive cells (**Figure 3C**). In contrast, cells expressing cGAS-rs200818241 exhibited an extended replicative lifespan (**Figure 3A**) accompanied by reduced activation of the canonical cGAS-STING pathway and p16 expression (**Figure 3B**), and a reduced number of SA-β-gal positive cells (**Figure 3C**) compared to cells expressing cGAS-WT. These findings indicate that rs200818241 protects against cellular ageing phenotypes.

**Figure 3.**
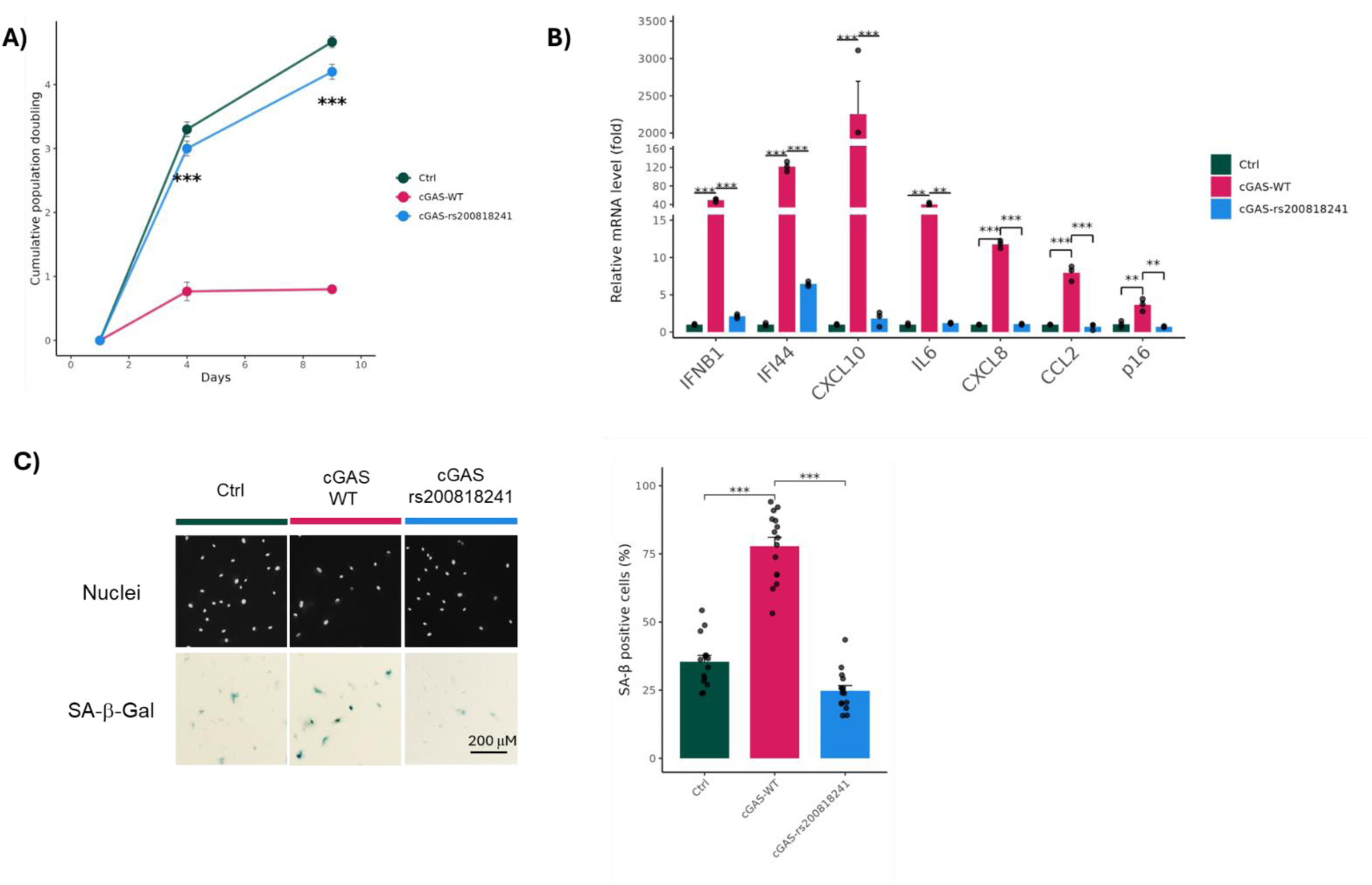
Longevity-associated cGAS variant rs200818241 delays cellular senescence in human mesenchymal stromal cells (hMSCs). **A**) Population doubling curves of cells expressing wild-type cGAS (cGAS-WT), the cGAS mutant (cGAS-rs200818241), or an empty vector control (Ctrl) in human mesenchymal stromal cells (hMSCs). Each dot indicates the cumulative population doubling per passage. cGAS-rs200818241 significantly extends cellular lifespan compared to cGAS-WT. **B**) qPCR analysis of downstream genes in the canonical cGAS-STING signaling pathway and the senescence-associated p16 gene (*CDKN2A*). Expression of cGAS-rs200818241 increases both downstream genes in the canonical cGAS-STING pathway and p16 levels, while cGAS-rs200818241 reduces the expression of all these genes compared to cGAS-WT. **C**), Senescence-associated β-galactosidase (SA-β-gal) staining assay showing that cGAS-WT increases the proportion of β-gal-positive cells, whereas cGAS-rs200818241 decreases the number of β-gal-positive cells compared to cGAS-WT. The left panel shows representative images of nuclei and SA-β-gal staining, the right panel presents the statistical analysis of β-gal-positive cells. All experiments were performed in triplicate. Statistical significance is denoted as follows: *p<0.05, **p < 0.01, ***p < 0.001.

### The observed cellular effects of rs200818241 in cGAS are conserved between humans and mice

The rs200818241 variant is conserved in mice (**Extended data 5C**). To assess whether the cellular effects of its functions are conserved across species, we created AN3-12 mouse embryonic stem cells (mESCs), harbouring the variant in a homozygous state. We subsequently determined the effect of the variant on cGAS-Sting activity using western blotting, both under normal conditions and after stimulating the mESCs with foreign double stranded DNA to activate the pathway. The unstimulated cGAS-rs200818241 mESCs showed a significant reduction in cGAS protein levels, which became even more apparent after stimulation of the cells (**Figure 4A**). Moreover, the cGAS-WT mESCs showed a significant downregulation of Sting, which is indicative of its activation [36], and an increased phosphorylation of Tbk1 after stimulation, while the cGAS-rs200818241 mESCs remained unaffected (**Figure 4B-C**). These findings indicate that the dampened cellular responses of rs200818241 upon stimulation are conserved between human and mice, opening the door for future experiments in model organisms to assess the physiological effect of this variant.

**Figure 4.**
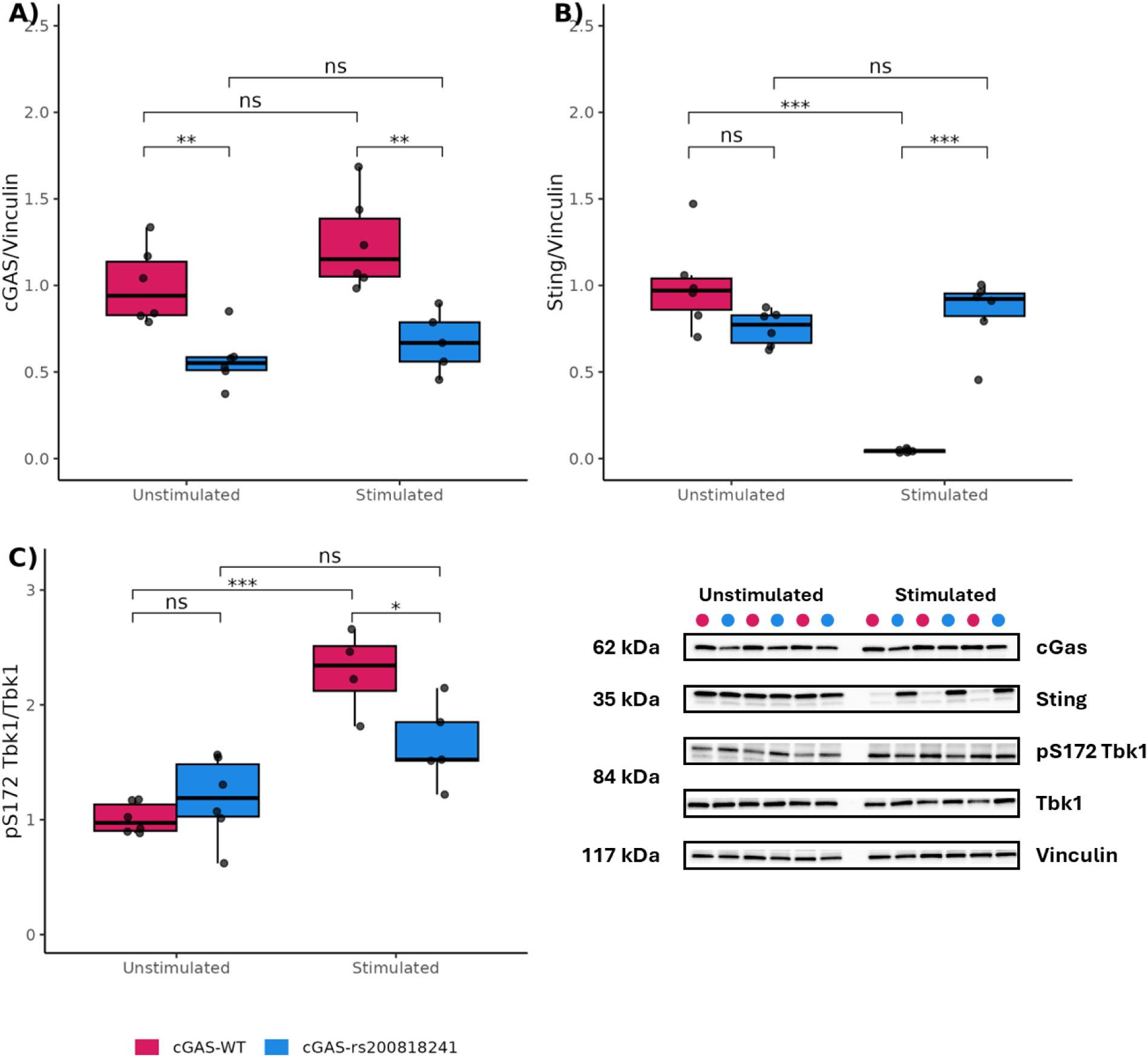
rs200818241 affects cGas-Sting activity in mouse embryonic stem cells (mESCs) **A**) Significant reduced response of cGas in cGAS-rs200818241 mESCs compared to cGAS-WT mESCs upon stimulation by transfection for 24 hours. **B**) Significant reduction of Sting in the cGAS-WT mESCs, but not the cGAS-rs200818241 mESCs, after stimulation by transfection for 24 hours. **C**) Significant increase in phosphorylated Tbk1 (S172) over total Tbk1 in the cGAS-WT mESCs, but not the cGAS-rs200818241 mESCs, after stimulation by transfection for 24 hours. Vinculin was used for normalisation. The data shown is from two independent experiments with three technical replicates each. Data was analysed using a one-way ANOVA and Dunnett’s post hoc test. Statistical significance is denoted as follows: ns; not significant,*P < 0.05, **P < 0.01, *** p < 0.001.

## Discussion

Long-lived families have a remarkable health and survival advantage across the life course [7, 11, 13]. However, the genetic architecture of longevity remains largely unresolved, with few common variants identified, suggesting a significant role for rare variants [18, 19]. Here, we used our recently established criteria [8] to select for nonagenarian sibships specifically enriched for ancestral longevity to identify novel longevity loci through hypothesis-free genetic linkage analysis. We identified four significant linkage regions (*Chr1q21.1, Chr6p24.3, Chr6q14.3, and Chr19p13.3*), which we followed up by sequence analysis of those regions using a filtering and validation approach, prioritizing protein-altering rare variants meeting strict criteria. We identified 12 high impact protein-altering rare variants in seven genes (*NUP210L, SLC27A3, CD1A, CGAS, IBTK, RARS2*, and *SH2D3A*), that we propose as novel longevity-candidate variants. One of these is rs200818241, present in two independent long-lived families. rs200818241 alters the function of cGAS *in vitro* in both human and mouse cells, hinting that this mutation protects against cellular ageing phenotypes.

Relative to community-dwelling nonagenarians of a comparable birth cohort, long-lived LLS nonagenarian sibships fulfilling the ancestry-based criteria scored significantly better on lifespan, cognition, and well-being, but not on functional status. These observations confirm prior studies showing that long-lived families have an improved memory performance and increased survival [37–39]. Our results highlight how using an ancestry family-based definition of longevity can identify individuals who will reach an extended lifespan and can help to identify health patterns associated with familial longevity. Further research should be undertaken to determine the causal direction of our findings.

The peak regions identified in the linkage analysis harbour genes previously found to be associated with ageing in candidate-based studies, including: *LMNA* [40], *IL6R* [41], *CRP* [42], *INSR* [43], *SHC1* [44], and the innate immune sensor for intracellular DNA and activator of STING signaling: *IFI16* [45]. We did not observe rare variants in these candidate genes that met all the selection criteria. The seven genes we identified with high impact protein-altering mutations may be involved in influencing longevity by processes that are increasingly altered with ageing, e.g. cancer, inflammation, cellular senescence or mitochondrial dysfunction [43] (for more information on their gene functions *see* ***Supplementary Table 10***). For two genes we identified a single rare missense variant in two independent LLS families: *SH2D3A* and *CGAS*. *SH2D3A* is a paralog of breast cancer anti-estrogen resistance gene 3 (*BCAR3*/*AND-34*) which plays an important role in the development of estrogen resistance in the progression of breast cancer [46]. Increased *SH2D3A* expression levels have been found to be associated with detrimental outcomes in cervical-, lung-, and pancreas-cancer. [47–49]. The major link to known pathways of biological ageing is represented by the variant (rs200818241) in *CGAS*, which was previously implicated in age-related inflammation and cellular senescence [26–28, 42]. This rare missense variant was selected for functional follow up studies. rs200818241 was more frequently present in German LLI centenarians compared to non-long-lived controls, suggesting that the variant may contribute to longevity more broadly outside of the familial context. *CGAS* encodes for a cytosolic DNA sensor that activates the innate immune system via the canonical cGAS-STING pathway leading to the activation and production of type I interferons, interferon stimulated genes, and pro-inflammatory cytokines [34]. *CGAS* and its homologs are broadly conserved across both multicellular and single-celled organisms [50]. Overactivation of the canonical cGAS-STING pathway has been associated with detrimental effects on tissue homeostasis and ageing. Inhibition of STING has been proposed as a strategy to slow age-related degeneration [30]. Yet studies of human genetic variation in *CGAS* remains sparse [51]. To our knowledge, no potentially protective human variants in this gene have previously been implicated in longevity. In naked mole-rats (*Heterocephalus glaber*), however, four C-terminal mutations (S463, E511, Y527, T530) in cGAS negated its inhibitory effect on homologous recombination repair, enhancing genome stability, delaying organismal aging and extending lifespan and healthspan [52]. Notably, these mutations are located in the region and in proximity of which variant rs200818241 (D452V) resides in the human cGAS protein.

Our functional assays suggests that rs200818241 acts as a reduced-function variant. It reduces cGAS protein stability and accelerates its turnover, thereby tempering cGAS-STING activation without abolishing activity. Such modulation may allow tighter control of innate immune responses and limit chronic inflammation [53]. We further explored the role of rs200818241 in cellular senescence, which is a state of irreversible cell cycle arrest induced by various environmental and cellular stressors, and is known to serve as a critical barrier for tumour formation. Senescent cells, characterized by a pro-inflammatory secretory phenotype, accumulate with age and are a major driver of ageing and ageing-related pathologies [54]. Previous studies have demonstrated that cGAS plays a key role in the induction of cellular senescence [28, 29]. Consistent with this, rs200818241 delayed proliferation arrest, reduced p16 expression, and decreased senescence-associated ß-galactosidase activity compared to the wildtype. These findings show how rs200818241 ameliorates cellular senescence likely through increased cGAS protein turnover and reduced signalling intensity. The effect was conserved in mice, where the variant elicited a similarly dampened cGAS-STING response, supporting a shared biological mechanism and providing rationale for future *in vivo* studies of this variant. cGAS inhibitors (small molecules such as VENT-03, JO14, XL-3156, cyclopeptide and flavonoid-based compounds) are being developed to treat autoimmune and autoinflammatory diseases [55]. However, while dampening the canonical cGAS-STING pathway may support inhibiting an overactive immune response, boosting this pathway has been pursued as a cancer therapy [59], reflecting its context-dependent roles in different physiological settings. Model organisms, such as mice or fish, could help assess the physiological impact of this variant on healthspan and lifespan. Similar functional approaches could be extended to the other rare variants identified in our linkage study.

This study comes with several limitations. We lacked a sufficiently large control population sequenced on the same platform, which prevented us to perform a well-powered genetic burden analysis to assess the enrichment of rare variants in the linkage regions. To address this, we implemented a rare variant prioritization strategy to identify likely (familial) longevity candidate variants with a predicted high impact on the genes. Moreover, while the functional assays were limited to cell-based *in vitro* assays, we conducted experiments across multiple cell lines from humans and mice and observed that its effects were conserved across species. Finally, our findings were derived from individuals of Northern-European descent. To strengthen the generalizability of the results, future research should include individuals from broader and more diverse populations. This may validate the role of cGAS in ageing and (familial) longevity and the impact of its rare variants across different genetic backgrounds and environments.

In summary, we characterized the *in vitro* functional impact of the rare missense *CGAS* variant rs200818241, which was identified in one out of four chromosomal regions linked to familial longevity. The *CGAS* variant, dampening the canonical cGAS-STING pathway and cellular senescence, was shared across two independent long-lived families contributing to the linkage. In addition, we identified variants in six other candidate genes involved in (familial) longevity warranting further investigation. Our study highlights the canonical cGAS–STING pathway as a potential mediator of protective organism-wide anti-ageing processes.

## Extended Data

**Extended data 1.**
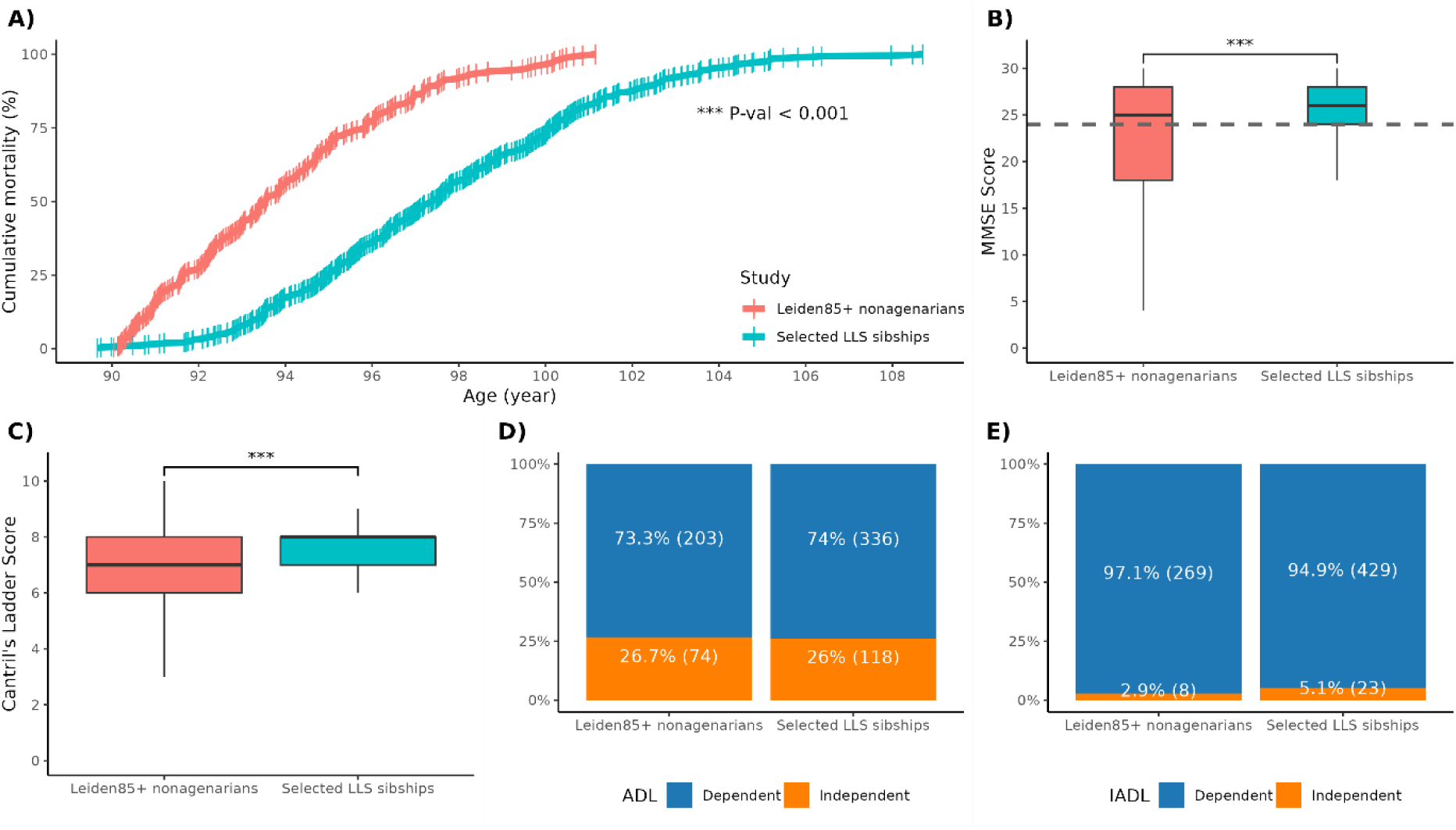
Comparison of selected LLS sibships with the general community-dwelling nonagenarian population from the Leiden 85+ study on common geriatric measurements. Additional details are provided in **Supplementary Table 1. A**) Cumulative mortality from age of inclusion till death. Red upper line show the Leiden 85+ nonagenarians (N=272; mean age of inclusion=90), blue lower line indicates the selected LLS sibships (N=479; mean age of inclusion= 93.6), **B**) Mini-Mental State exam (MMSE)-score for the Leiden85+ nonagenarians (N=277) and selected LLS sibships (N=431). The grey dashed line (MMSE=24) indicates the screening cutoff score for mild-cognitive impairment. **C**) Quality of life Cantril’s Ladder scoring for the Leiden85+ nonagenarians (N=201) and selected LLS sibships (N=417). **D**) Activities of Daily Living (ADL) dependency in for the Leiden 85+ nonagenarians (N=277) and selected LLS sibships (N=452). **E**) Instrumental Activities of Daily Living (IADL) dependency for the Leiden 85+ nonagenarians (N=277) and selected LLS sibships (N=452). For untransformed distribution of the (I)ADL scores see **Extended Data 2.**

**Extended data 2.**
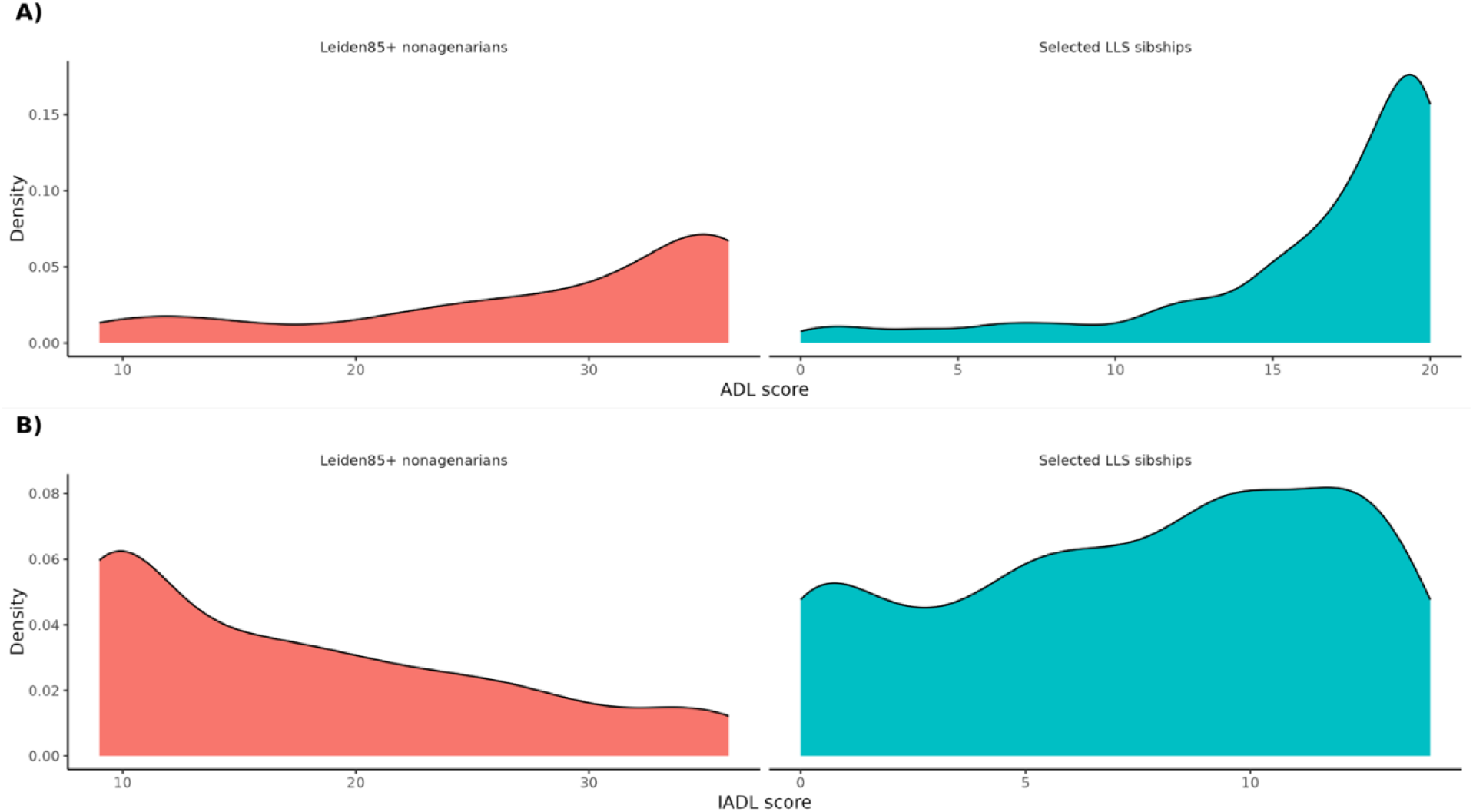
ADL & IADL-scoring distribution Leiden 85+ nonagenarians and Selected LLS Sibships. **A**) Distribution of the Activities of Daily living (ADL)-scoring to asses a person’s ability to perform basic self-care tasks needed for independent living. Within the Selected LLS Sibships (N=454) scoring was done using Katz-ADL (0-20). In the Leiden85+ nonagenarians (N=277) scoring was done using the Groningen Activity Restriction scale (GARS)- ADL component (9-36), the polarity from the original score was inverted to ensure that in both groups a higher score means more functionally independent (better). **B**) Distribution of the Instrumental Activities of Daily Living (IADL)-score which asses a person’s ability to live independently in the community by managing more complex everyday tasks. Within the Selected LLS Sibships (N=452) scoring was done using Lawton’s-IADL (0-14). In the Leiden85+ nonagenarians (N=277) scoring was done using the GARS IADL component (9-36), the polarity from the original score was inverted to ensure that in both groups a higher score means more functionally independent (better).

**Extended data 3.**
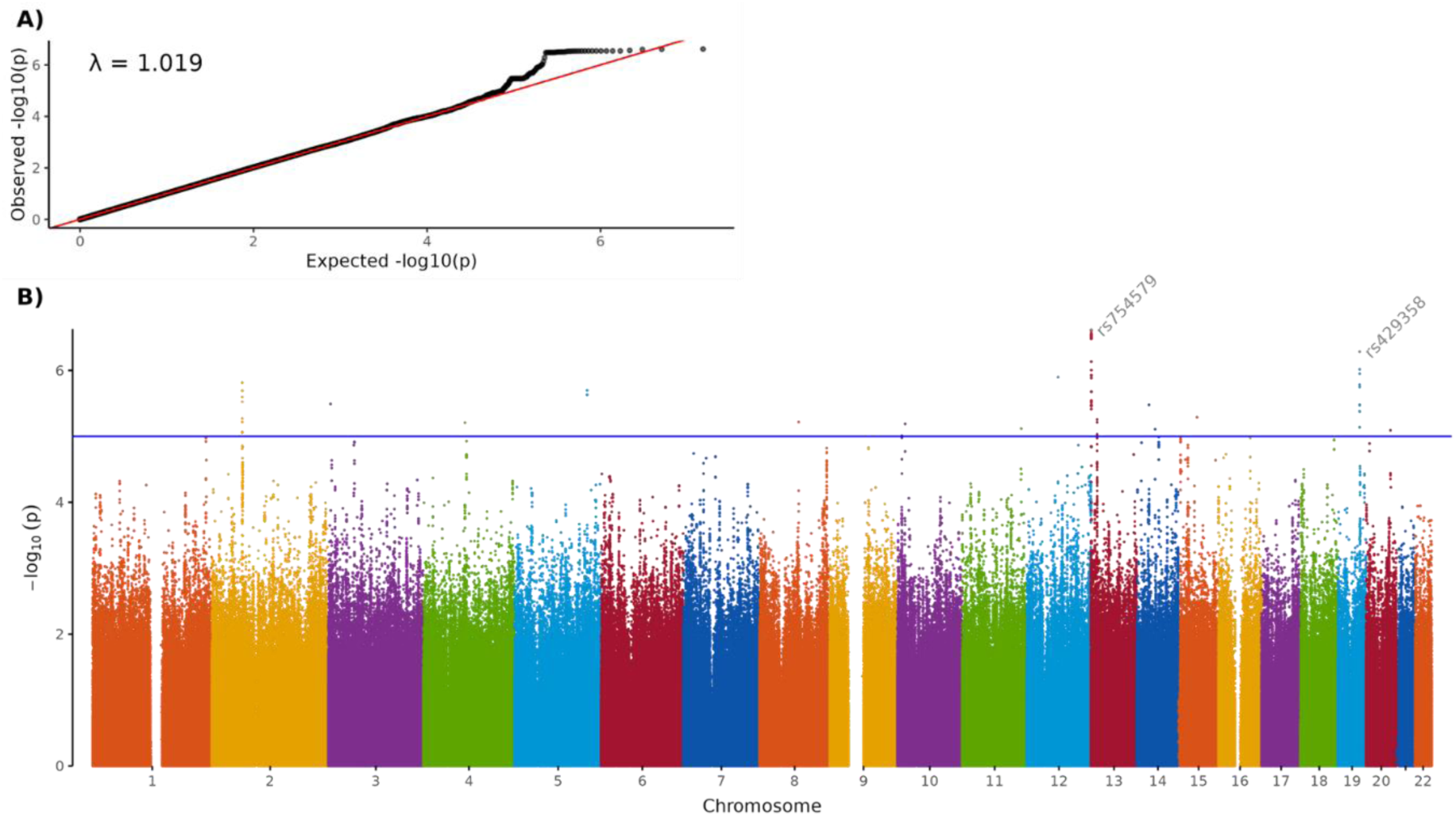
Genome-wide association analysis (GWAS) of familial longevity within the Leiden Longevity Study (LLS). This GWAS utilizes HRC v1.1 imputed SNP genotype microarray data of 472 long-lived nonagenarian siblings coming from 212 selected LLS sibships enriched for familial longevity, and 473 general population partners as controls exhibiting no familial longevity from the LLS. The association was performed using SAIGE [56], using sex as a covariate and corrected for sample relatedness. **A).** Quantile-quantile plot for association summary statistics from single-variant analysis. **B).** Manhattan plot of association summary statistics from single-variant analysis. The blue line indicates the genome wide suggestive line of P-value < 1·10^-5^. Variants labelled with the rsID are the tophits that passed a P-value < 1·10^-6^. No variants passed the genome wide significant threshold of P-value < 5**·**10^-8^, and we did not identify any suggestive SNP hits within our four linkage regions. For detailed results see **Supplementary Table 4**.

**Extended data 4.**
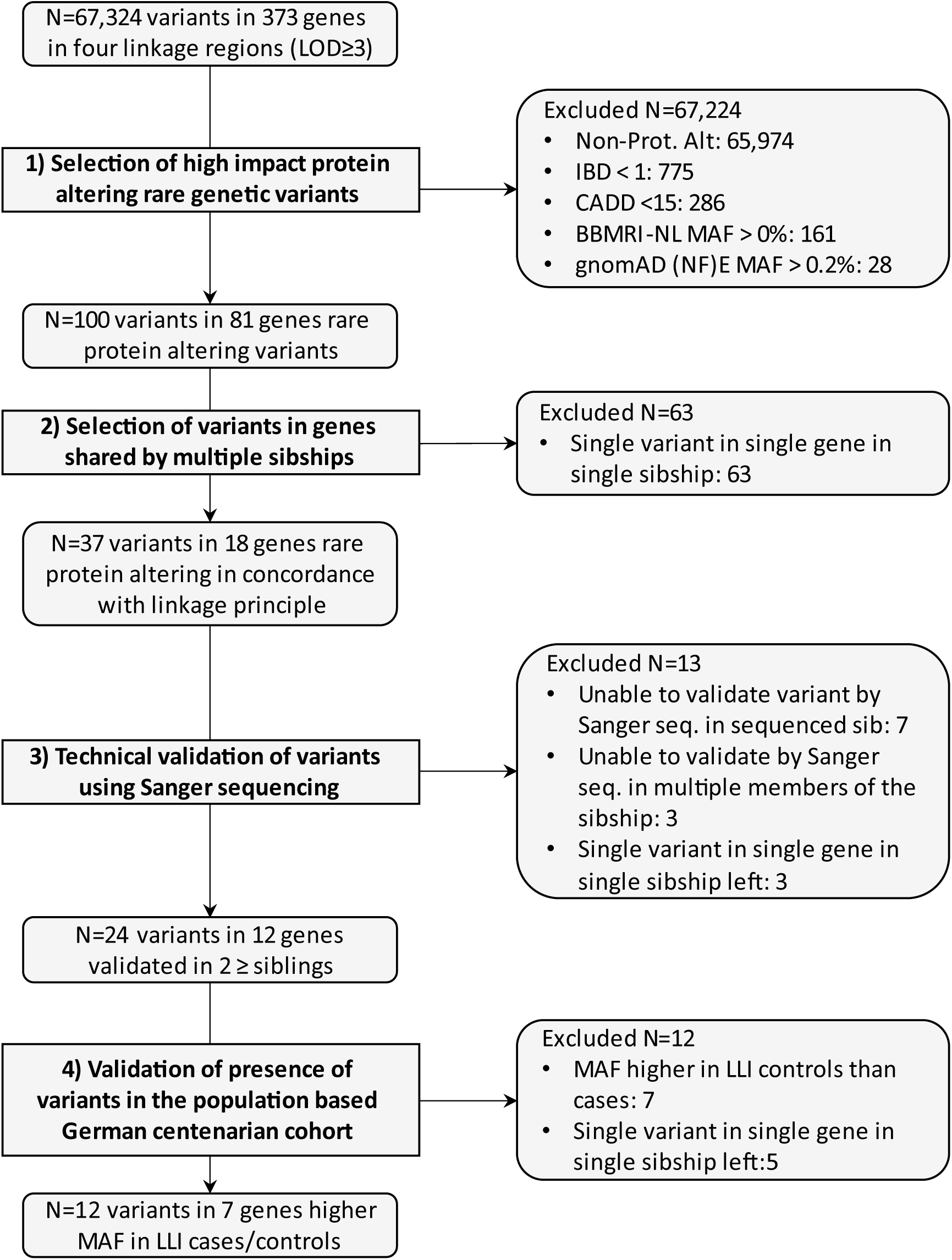
Flowchart of filtering and validation steps of genetic variants observed in the linked regions carried by the individuals in the selected LLS sibships. **Non-Prot. Alt:** Non-protein-altering, protein-altering: missense, start lost, stop gained stop lost; **IBD**: Identity-by-Descent; **CADD**: Combined Annotation Dependant Depletion score; **BBMRI-NL MAF**: Dutch population Minor Allele Frequency (MAF); **gnomAD (NF)E MAF**: Genome Aggregation Database (gnomAD) Non-Finnish (NF) European MAF; **Sanger seq.**: Sanger sequencing; **LLI MAF**: Long-Lived individuals (LLI) German population MAF, Cases: ≥94 years (including 643 centenarians), Controls: younger population (mean age 68 years).

**Extended data 5.**
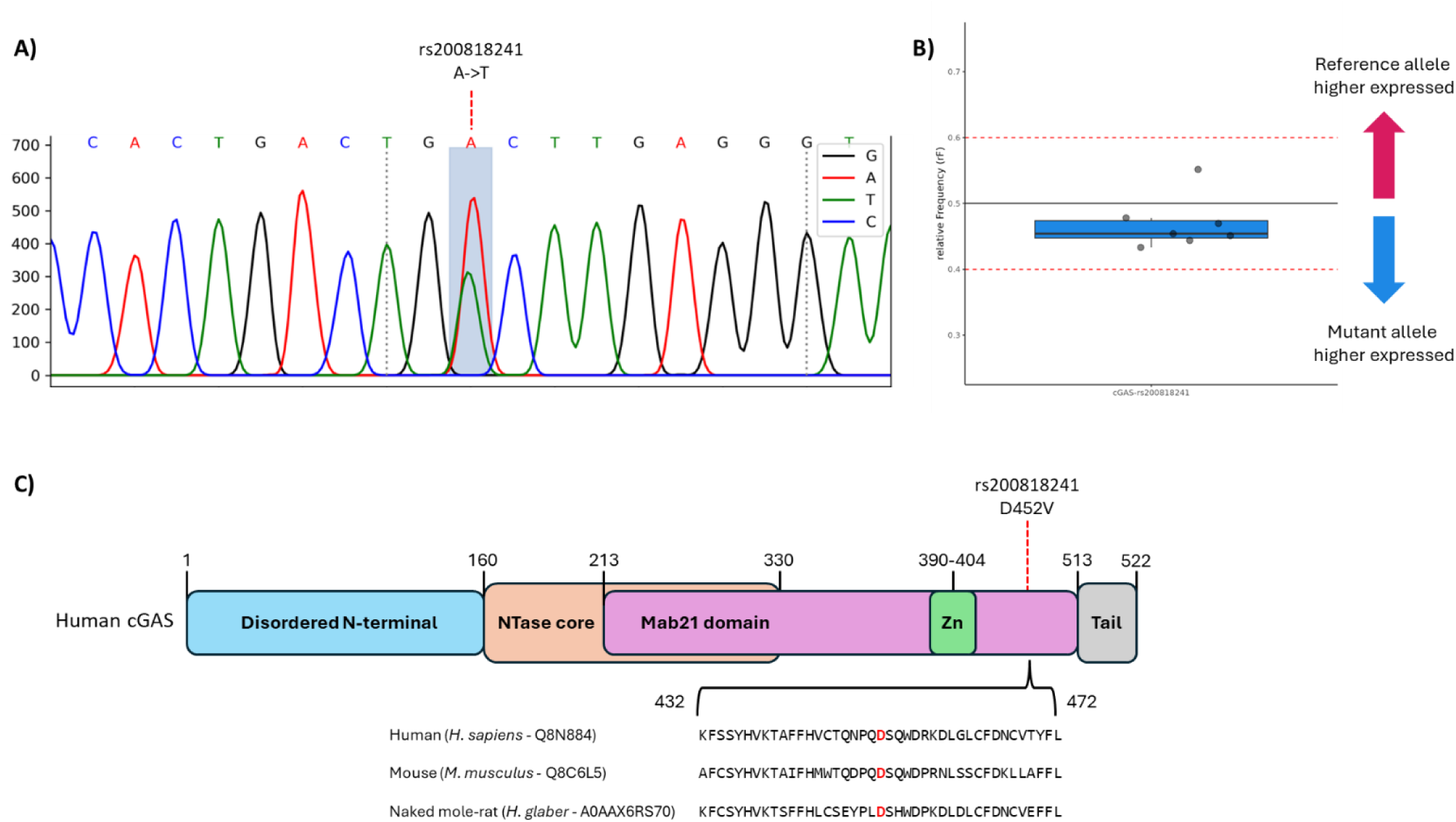
Location, allelic-specific expression (ASE), and conservation of the rare missense cGAS variant rs200818241. **A)** Sanger sequencing validation of rs200818241 in a representative carrier from one of the identified families, with the variant highlighted in blue. **B)** ASE analysis of rs200818241–cGAS variant using a TaqMan® genotyping assay in F2-selected LLS sibships and their F3-offspring variant carriers. No evidence of ASE was observed in any of the carriers. **C)** Schematic representation of the cGAS protein structure showing functional domains and the position of variant rs200818241 (indicated by a red dashed line). Sequence alignment of cGAS from human (UniProtKB: Q8N884), mouse (UniProtKB: Q8C6L5), and naked mole-rat (UniProtKB: A0AAX6RS70) is shown below, with the residue corresponding to the rs200818241 site highlighted in red.

**Extended Data 6.**
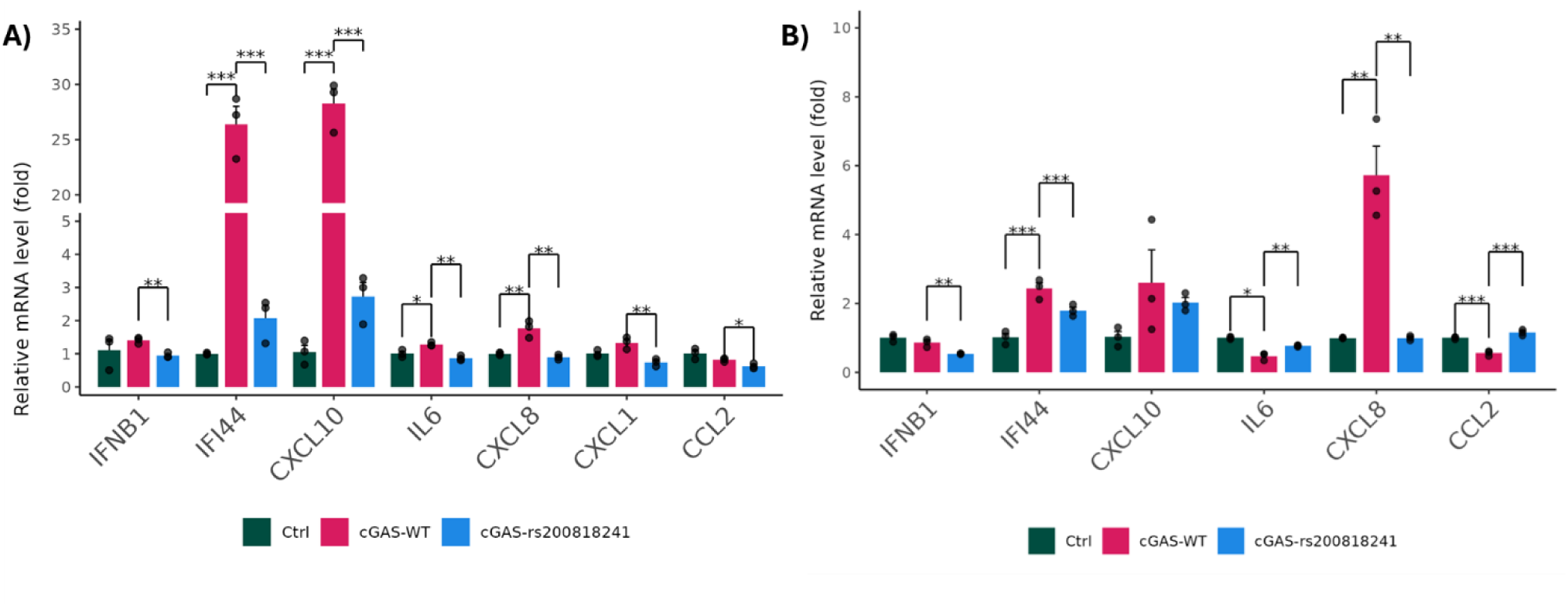
Longevity-associated cGAS variant suppresses the activation of cGAS-STING pathway in a cell type-specific manner. **A**) qPCR analysis of cGAS-STING downstream gene expression in primary human astrocytes expressing wild-type cGAS (cGAS-WT), the cGAS mutant (cGAS-rs200818241), or an empty vector control (Ctrl). The cGAS-rs200818241 significantly suppresses the activation of cGAS-STING downstream genes. **B**) qPCR analysis of downstream gene expression in human fibroblasts (IMR-90) expressing cGAS-WT, cGAS-rs200818241, or Ctrl. In fibroblasts, the cGAS-rs200818241 only suppresses a subset of cGAS-STING downstream genes, including *IFNB1*, *IFI44*, and *CXCL8*. All experiments were performed in triplicate. Statistical significance is denoted as follows: *p<0.05, **p < 0.01, ***p < 0.001.

## Methods

### Study populations

#### Leiden Longevity Study

In this study, we investigated a subset of the population included in the Leiden Longevity Study (LLS) [9]. The LLS was initiated to study the genetic determinants of human longevity, and is a family-based study consisting of 421 independent nonagenarian proband sibships (F2, N=944) of Dutch European descent, with a mean age at inclusion of 93.4 years (89-103), their proband offspring (F3, N=1671) (59.4 years, 34-80), and the offspring partners (F3, N=744) (58.7 years, 30-79). In addition, parents (F1 parents) were linked to the nonagenarian sibships using genealogical records, and mortality information of the parental generation of the F3 partners was collected. The proband sibships were recruited between 2002 and 2006. Families in the LLS were included if at least two siblings met the following criteria: 1) two long-lived siblings were alive and willing to participate, 2) they met the age criteria with the men being at least 89 years old and women at least 91 years old. 3) the proband sibships had to have the same mother and father and being of Dutch European descent. This age category of the proband sibships represents <0.5% of the Dutch population in 2001.

The Leiden Longevity Study protocol was approved by the Medical Ethical Committee of the Leiden University Medical Center before the start of the study (P01.113) on the 16^th^ of August 2002. The first participant was enrolled on the 5^th^ of September 2002. In accordance with the Declaration of Helsinki, the Leiden Longevity Study obtained informed consent from all participants prior to their entering the study.

#### Leiden 85+ study

The Leiden 85+ study is a prospective population-based study in 85-year-old inhabitants of the city of Leiden, the Netherlands [23] . The Leiden 85+ study encompasses participants (N=599) from the 1912-1914 birth cohorts living in the city of Leiden that were recruited between September 1997 and September 1999. No exclusion criteria were used. Participants were visited at baseline, and yearly up to the age of 90 years, at their place of residence. Mortality information is available (N=594) until February 2014, with five participants still alive at the time of censoring.

The Leiden 85+ study protocol was approved by the Medical Ethical Committee of the Leiden University Medical Center (P097.04). Informed consent was provided by the participants or, in case of severe cognitive impairment, by the most significant other .

#### German long-lived individuals (LLI) study

The German LLI cohort comprises 1,265 samples of unrelated LLI (mean age: 99 years, age range: 94–110 years), including 602 centenarians (≥100 years), and 4,195 younger population-representative controls (age range: 18–83 years, mean age: 50.5 years). The male/female ratio in the sample was 1:2.8. All participants were of German ancestry. LLI showed no overt signs of major diseases or cognitive impairment [19].

Written informed consent to participate in the study was obtained from all participants, and the project was approved by the Ethics Committee of the Medical Faculty of Kiel University.

#### Subsampling of sibships enriched for ancestral longevity within the Leiden Longevity Study and nonagenarians individuals from the Leiden 85+ study

In order to maximize the enrichment for cases with the genetic longevity component we applied a more stringent inclusion criterium of the nonagenarian siblings pairs of the original LLS population [22]. This LLS subset is referred to as selected LLS sibships. We applied the following additional inclusion criteria: 1) a minimum of one parent (F1) belonging to the top 10% survivors of his/her birth cohort and 2) at least two nonagenarian siblings (F2) in a sibship belonging to the top 10% survivors of their respective birth cohort.

We used population-based lifetables from the Netherlands to calculate birth cohort and sex-specific percentiles for each individuals in the LLS proband generation, and their parents [57]. This approach prevents against the effect of secular mortality trends of the last centuries and enables comparisons across study populations [4, 58]. These lifetables contained, for each birth year and sex, an estimate of the hazard of dying between ages x and x + n (hx) based on yearly intervals (n = 1) up to 99 years of age. Conditional cumulative hazards (Hx) and survival probabilities (Sx) were derived using these hazards. In turn, we could determine the sex and birth year specific survival percentile for each person in our study. For instance, a female born in 1910 who lived to age 92 belonged to the top 10% of her birth cohort, based on Dutch lifetable information, meaning only 10% of women born in 1910 lived to an older age. In contrast, a woman born in 1850 who also lived to age 92 would have been in the top 2% of her birth cohort. 212 (50.4%, N=479) selected LLS sibships met the criteria for enrichment for ancestral longevity of the original 421 enrolled nonagenarians sibships (**Table 1**).

Within the Leiden 85+ study we selected participants who reached at least 90 years of age, regardless of their ancestral longevity, and did partake in the final examination round in the study. We call these the Leiden 85+ nonagenarians (N=277 out of 599 85+ individuals in total), and use them to compare with the selected LLS sibships on common geriatric assessments. As both populations are of similar birth-cohort, age, cultural-, and genetic background (**Supplementary Table 1**).

#### Geriatric assessment of the nonagenarians in the selected LLS sibships and Leiden 85+ subsample

In both the LLS and Leiden 85+ study global cognitive function was measured with the Mini-Mental State Examination (MMSE) [59] and general subjective well-being was assessed using Cantril’s Ladder of Life [60]. MMSE scores range from 0 point (severe cognitive impairment) to 30 points (optimal cognitive function). Cantril’s Ladder of Life range with a score range from 1 (worst) to 10 (optimal) self-perceived well-being. Within the LLS, functional status was assessed by Katz Activities of Daily Living (ADL) [61] and Lawton’s Instrumental Activities of Daily living (IADL) [62]. ADL disability scores range from 0 (fully dependent in all activities) to 20 points (fully independent in all activities). IADL disability scores go from 0 points (fully dependent in all activities) to 14 points (fully independent in all activities). In the Leiden 85+ study functional status was questioned through the Groningen Activity Restriction Scale (GARS) [63] and split in both ADL and IADL components. The polarity from the GARS-scores was inverted, ranging from 9 (fully dependent in all activities) to 36 points (optimal) to ensure that in both groups a higher score means more functionally independent. For our comparative analysis we dichotomized IADL and ADL scores into 0 (cannot or only with help from others) or 1 (fully independently, with or without difficulty). The dichotomizing of the score was done because no universal cut-off points of the GARS are known [64]. See **Extended data 2** for the distributions of the untransformed scores.

#### DNA Isolation – Genotyping LLS

DNA from the LLS was extracted from buffy coat samples at baseline using conventional methods [65]. Cases were genotyped using Illumina Infinium HD Human660W-Quad Beadchips (Illumina, San Diego, USA). Quality control included SNP-wise call rate (≥95%), their minor allele frequency (>1%) and derivation from the Hardy–Weinberg equilibrium (p < 1 x 10^-4^) [66]. SNPs were imputed against the Haplotype Reference Consortium (HRC) v1.1 reference panel.

#### Affected sib-pair linkage analysis

Affected sib-pair (ASP) non-parametric linkage (NPL) analysis, and error correction was carried out using the Multipoint Engine for Rapid Likelihood Inference (Merlin) v1.1.2 software package on the selected LLS sibships [67]. From the genotype microarray data unlikely recombinants were detected and removed with the -pedwipe module in Merlin. Additionally, Pearson’s r^2^ was estimated using the total data set. All SNPs had a pairwise r^2^ <0.4, for a total genome-wide SNP set of 11,926.

A non-parametric Kong & Cox allele-sharing LOD-score [68] was computed using the -npl module in Merlin with the ALL scoring function, that uses a linear model to evaluate linkage.

This model allows to detect “small” increases in allele sharing spread across a large number of families, which is what one usually expects in a complex disease [67].

A peak maximum (LOD_max_) score of ≥3 was considered as significant and a LOD of >2 as suggestive for linkage, which corresponds to a genome-wide (chi-squared) P-value of 2.0·10^-4^ and 2.4·10^-3^, respectively [69]. By applying this moderate cutoff, instead of focusing on the more extreme LOD>3.3 (chi-square P-value = 9.8·10^-5^) and the 1-LOD drop, allowed us to explore biological evidence for the linkage in a maximal number of genes and variants. Linkage regions linked with familial longevity were delimited by SNPs in the LOD_max_ with a LOD-score >2.

#### Genome-wide association analysis in the LLS

Genome-wide association analysis (GWAS) of familial longevity was performed using imputed genotype microarray data from the LLS. The analysis was conducted using Scalable and Accurate Implementation of Generalized mixed model (SAIGE) [56], to account for case-control imbalance and sample relatedness. To maximize the contrast between groups, the longevity phenotype was dichotomized. Cases comprised the 472 (of the 479; 98.5%), of the selected long-lived nonagenarian siblings, while controls included 473 unrelated general population partners from the LLS who did not exhibit familial longevity. The analysis was adjusted for sex and relatedness to control for potential confounding effects.

#### Whole-genome sequencing data in the Leiden Longevity Study

Whole-genome Sequencing (WGS) was generated between 2011-2013, in LLS nonagenarians (F2) that 1) had DNA available and blood available of their offspring (F3), 2) had known lifespan of their parents (F1), and 3) had to score low on the Family Mortality History Score (<= -1.05) [70], as evidence that the mortality hazard in the parents is lower than their birth cohort. If multiple siblings within a sibship met these criteria, the oldest or the one still alive was selected from these families for WGS [25]. Resulting in 179 selected LLS sibships for which WGS data was available in one sibling.

WGS data generation in the subset of LLS nonagenarians was performed by Complete Genomics (Complete Genomics Inc., Mountain View California version 1.3.0) at a coverage of >30x. Quality control was performed on the data using the flags generated by the Complete Genomics platform. Variants with VQLOW, FET30 or the AMBIGUOS flag were filtered out; VQLOW: generally fewer than seven reads, FET30: somatic variants with a Fisher somatic score of ≤ 30 (corresponding to a p-value < 0,001), AMBIGUOUS: alleles where there is strong evidence that the allele is not reference, but with insufficient evidence to make a single high-confidence call.

#### Private Rare Variant Filter and Validation pipeline

To identify rare predicted protein-altering high impact variants in the positive linkage regions that may contribute to the linkage and allow for follow up functional genomics analyses, a collapsing filtering strategy was applied (**Extended data figure 4**). Variants in the linkage regions, including single nucleotide variants (SNVs) and indels, were annotated using the Ensembl Variant Effect Predictor (VEP) [71], using the Ensembl genome assembly GRCH37 build 92 as reference assembly. VEP provides information on gene annotation, amino acid change annotation and dbSNP ids (build 150) [72].

1. **Selection of high impact protein-altering rare genetic variants** VEP annotated variants were filtered for protein-altering consequences (missense, start_lost, stop_gained_stop_lost). To increase the probability that siblings carry the same selected variants allele sharing, IBD, was calculated using the –ibd module from Merlin [67] and variants were selected to be IBD ≥ 1. CADD v1.3 was used to measure the deleteriousness and high impact of the variants, a CADD score > 15 was defined as deleterious variants [73]. To verify that the identified deleterious protein-altering variants are rare, we utilized sequencing data assayed on 100 individuals of Dutch Caucasian origin (≤65 years old) which was collected by the Dutch Biobanking and Biomolecular Resources Research Infrastructure initiative (BBMRI-NL) [74, 75]. Participants of BBMRI-NL are not selected for particular characteristics other than that they should reflect a random sample of the apparently healthy Dutch population. Variants had to be absent in BBMRI-NL (MAF=0%). Next, the rarity of the variants were filtered against the (Non-Finnish) European (NFE) general population in the Genome Aggregation Database (gnomAD) v2.1.1 database [26], variants were considered rare if they occurred with a NFE-MAF of <0.2%.
2. **Selection of variants in genes shared by multiple sibships** In order to enhance the likelihood that the selected high impact protein-altering rare genetic variants are contributing to the genetic linkage, and be associated with familial longevity, variants had to be either: 1) present in multiple sibships, or 2) multiple variants were detected in the same gene across at least two sibships
3. **Technical validation of variants using Sanger Sequencing** Sanger sequencing was applied to validate whether the sibships were actually carrying the identified variant. As such primers were designed flanking all selected variants (**Supplementary Table 5**), using the NCBI PRIMER-BLAST tool [76]. PCR products were purified using a AMPure XP Reagent Kit (Beckman Coulter Life Sciences, Indiana, United States) and sequenced using an automated ABI 3730XL sequencer (Applied Biosystems, Forster City, United States). Steps taken to technically validate the variant where as follows: 1) the DNA of the person in which the variant was identified was Sanger-sequenced for carrying the variant. 2) all members of the accompanying sibship that had DNA available were Sanger-sequenced for the identified variant. Only if the variant in the sequenced individual was replicated and multiple members of the sibship (≤2) were carriers, we considered it validated (**Supplementary Table 6**).
4. **Validation of presence of variants in the population based German centenarian cohort** For the purpose of considering whether the variants are rare and associated with longevity (i.e. healthy ageing) the allele frequency was compared between long-lived cases and younger controls from the German LLI study, a population ethnically and geographically similar to that of the Dutch populace.

Whole Exome Sequencing (WES) and variant calling of the German LLI was performed using standard methods [19]. The validated Sanger-sequenced variants were replicated in the German LLI study. The variant location was converted from genome build GRch37 to Grch38, using the UCSC LiftOver Tool, and allele counts (AC) were scanned for presence in the LLI cases and LLI controls, and variants were filtered out if they were more frequent in the controls than cases.

#### Allelic Specific Expression of rs200818241

The positional candidate *cGAS*-rs200818241 variant (Table 3) was tested for allelic imbalance using TaqMan^®^ genotyping assay using probes designed by the manufacturer and carried out using manufacturer standard guidelines (ThermoFisher, Waltham, United States). From the long-lived LLS families were the variant was identified, DNA (EDTA buffy coats) and RNA (PAXgene®, BD Biosciences, Franklin Lakes, United States) was isolated in the selected LLS sibships (F2), their offspring (F3), and partners of the offspring as controls (F3). cDNA was synthesized from the RNA using Transcriptor First Strand cDNA Synthesis Kit (Roche, Basel Switzerland). Samples were measured sixfold, where 10ng gDNA and cDNA were loaded along with the TaqMan^®^ probes in 384-Well plates and genotyped and quantified using QuantStudio^™^ 6 Flex Real-Time PCR System (ThermoFisher).

### Functional assays for cell growth, protein stability, and senescence in human cell-lines

#### Cell culture

Human mesenchymal stem cells (hMSCs, ATCC) were cultured in Minimum Essential Medium alpha (MEM α) supplemented with 10% fetal bovine serum (FBS, Gibco), 1% penicillin/streptomycin (Gibco), and 1 ng/ml recombinant human basic fibroblast growth factor (bFGF, STEMCELL Technologies) on gelatin-coated culture dishes (Invitrogen). Human primary astrocytes (ScienCell) were maintained in astrocyt medium (ScienCell) on poly-L-lysine-coated plates (ScienCell), following the manufacturer’s protocol. Human KGN granulosa cells, were cultured in a 1:1 mixture of Dulbecco’s Modified Eagle Medium and Ham’s F-12 medium (DMEM/F-12, Gibco), supplemented with 10% FBS (GeminiBio) and 1% penicillin/streptomycin (Gibco). Human IMR-90 fibroblasts and HEK-293T cells were grown in DMEM (Gibco) supplemented with 10% FBS (GeminiBio) and 1% penicillin/streptomycin (Gibco). All cell lines were incubated at 37°C in a 5% CO₂ humidified incubator and regularly tested for mycoplasma contamination.

#### Virus preparation and cell transduction

The plasmid containing the cDNA of wild-type cGAS (pLenti-CMV-cGAS-HA) was obtained from Addgene. The cGAS variant (rs200818241) was introduced into the pLenti-CMV-cGAS-HA plasmid using the Q5® Site-Directed Mutagenesis Kit (New England Biolabs). To generate lentiviral particles, the cGAS plasmid, along with psPAX2 and pMD2.G (Addgene), were co-transfected into HEK-293T cells using Polyethylenimine (PEI, Sigma). Lentiviral supernatants were harvested and concentrated using the Lenti-X Concentrator (Takara Bio). For transduction, the concentrated lentivirus was applied to various human cell lines in the presence of 4 µg/ml polybrene (Sigma). Cells were harvested for analysis 24 hours post-transduction, or selected with 1 ng/ml puromycin for 48 hours to establish stable cell lines.

#### RT-qPCR

Total RNA was extracted from cells using the RNeasy Plus Kit (Qiagen) following the manufacturer’s protocol. One microgram of total RNA was reverse-transcribed into complementary DNA (cDNA) using the PrimeScript™ RT Reagent Kit (Takara). Quantitative real-time PCR (qPCR) was performed on the QuantStudio™ 6 Pro Real-Time PCR System (Applied Biosystems) with PowerUp™ SYBR™ Green Master Mix (Applied Biosystems). Data were normalized to GAPDH expression and analyzed using the ΔΔCq method. Primer sequences for all RT-qPCR assays using human cell-lines are listed in **Supplementary Table 8.**

#### SA-β-GAL staining assay

Cells were subjected to SA-β-GAL staining as previously described [77] . In brief, cultured cells were washed with PBS and fixed at room temperature for 4 minutes using a fixation buffer containing 2% formaldehyde and 0.2% glutaraldehyde. After fixation, cells were stained overnight at 37°C with freshly prepared staining solution to detect SA-β-galactosidase activity. The percentage of SA-β-GAL-positive cells was quantified and used for statistical analysis.

#### Western blotting

Cell lysates were prepared using Pierce™ Lane Marker Reducing Sample Buffer (ThermoFisher). Protein samples were denatured at 95°C for 3-5 minutes, followed by SDS-PAGE and electrophoresis. Western blotting was performed as previously described [78]. The following antibodies were obtained from Cell Signaling Technologies: anti-cGAS (#15102T), anti-STING (#13647T), anti-IRF-3 (#11904T), anti-p-STING (#50907T), anti-p-IRF-3 (#37829T), and anti-GAPDH (#32118S)

#### CHX chase assay

Cells were exposed to 50 µg/ml CHX for the specified durations, and protein degradation was assessed by Western blotting as previously described [78].

#### Growth curve assay

Cell population doubling was assessed as described previously [79]. In brief, the number of human mesenchymal stem cells (MSCs) was counted at each passage. The population doubling per passage was calculated using the formula log₂(number of cells collected / number of cells seeded). Cumulative population doublings were determined and plotted over time.

#### Generation and culturing of (transgenic) mESCs

AN3-12 mESCs harbouring cGAS rs200818241 were generated and cultured as previously described [20]. In short, we used the pSpCas9n(BB)-2A-GFP (PX461) vector (Addgene plasmid #48140), containing two guide RNAs (gRNAs) targeting the *cGAS* locus, and a 120 bp single-stranded DNA oligonucleotide containing the mutation of interest, as well as a silent mutation to disrupt the protospacer adjacent motif for gRNA2 (**Supplementary Table 9**). After transfecting the mESCs using Lipofectamine 3000 (Thermo Fisher Scientific, Massachusetts, United States), haploid and GFP-positive cells were single-cell-sorted using the BD FACSaria Fusion (BD Biosciences). After growing them for 10 days, the clone that showed a positive restriction digestion with DrdI (NEB) after PCR (details in **Supplementary Table 9**) was genotyped. The (transgenic) mESCs were maintained in a diploid state at 37°C with 5% CO2 in high glucose Dulbecco’s Modified Eagle Medium supplemented with 3.7 g/l Sodium bicarbonate, 13.7% FBS, 1% Pen-Strep, 1% L-Glutamate, 1% Sodium Pyruvate, 1% MEM Non-Essential amino acids, 0.1% beta-mercaptoethanol and 0.01% leukaemia inhibitory factor (Millipore).

#### Stimulation of the mESCs, protein extraction, and western blotting

To activate the cGas-Sting pathway, mESCs were transfected with 2 µg of the empty pSpCas9n(BB)-2A-GFP vector for 24 hours. Proteins were extracted using RIPA Buffer (Thermo Fisher Scientific) and subsequently used for western blotting with antibodies against cGas (#31659), Sting (#50494), Tbk1 (#3504), phospo-Tbk1 (Ser172) (#5483) (all from Cell Signaling), and Vinculin (#ab129002, Abcam). Details on the used procedures have been described previously [20].

#### Statistical analysis

Descriptive statistic were used to summarize the characteristics of the study groups.

The mortality analysis was adjusted for sex and performed using a left-censored Cox-type frailty (random effect) survival model to correct for late entry into the study set according to age. The mortality analysis was carried out using the FrailtyEM package [80]. A random effect (Gamma distribution) was used to adjust for similarity among family members. Families were defined by individuals sharing the same parents. A P-value ≤ 0.05 value was considered statistically significant. Statistical analyses were performed in R (v4.2.2) using base packages unless stated otherwise.

## Funding

The research leading to these results work was supported by the Netherlands Organization for Scientific Research (NWO; domain Health Research and Medical Sciences, 09120012010052) and the VOILA consortium (457001001). The Leiden Longevity Study received funding from the European Union Seventh Framework Programme (FP7/2007–2011; grant 259679). This study was supported by the Netherlands Consortium for Healthy Ageing (050-060-810), within the framework of the Netherlands Genomics Initiative, NWO, and BBMRI-NL (184.021.007 and 184.033.111). This project also received funding from the European Research Council (ERC) under the Horizon Europe programme (ElucidAge, 101041331). In addition, this work was supported by the United States National Institutes of Health (Y.S. AG056278 and AG017242). D.K received financial support from the Deutsche Forschungsgemeinschaft (DFG; RTG TransEvo 2501 – project 400993799).

Views and opinions expressed are those of the authors only and do not necessarily reflect those of the European Union or the ERC Executive Agency. Neither the EU nor the granting authority can be held responsible.

## Acknowledgements

The researchers would like to thank the study participants of the Leiden Longevity Study, the Leiden 85+ study, and the German Long-Lived Individuals study for their invaluable contributions. We thank Genomescan B.V. (Leiden, the Netherlands) and Michelle Bakker for their support with validation of the longevity candidate variants, Leon Mei for his bioinformatic input, and Ramona Pahl and Elina Singer for their help with the mESC experiments. We are grateful to Paul D. Robbins for generously providing the human KGN granulosa cells, and to Feng Zhang for gifting us the pSpCas9n(BB)-2A-GFP (PX461) vector.

## Contributions

P.C.P, M.B., P.E.S., and J.D. conceptualized the study. Y.S., P.E.S., and J.D. supervised the study. P.C.P., D.K., T.G., J.Y., and S.K. performed data analyses. P.C.P., D.G. H.H, N.L., H.E.D.S., G.C.L, E.B., A.A., and J.D. performed experimental analyses. S.T., N.M.A.B., M.B., and A.N. provided resources. P.C.P., P.E.S., and J.D. wrote the first draft of the manuscript. P.C.P., D.G., D.K., S.T., A.A., N.M.A.B., M.B., A.N., Y.S., P.E.S., and J.D. contributed to manuscript review and editing. All authors approved the final manuscript.

## Data availability

The LLS datasets analyzed in the current study are available following a data access procedure (https://leidenlangleven.nl/data-access/). For each request it will be tested whether the research is in compliance with the informed consent that has been signed by the LLS participants. All gel electrophoresis and microscopy data supporting this study will be included in the article and its supplementary information files.

## Code availability

This study does not report original code. All data were analysed and processed using published software packages, the details of which are provided and cited in the Methods section.

## Abbreviations

AC: Allele counts
ADL: Activities of Daily Living
ASP: Affected sib-pair
BBMRI-NL: Dutch Biobanking and Biomolecular Resources Research infrastructure initiative
bFGF: recombinant human basic fibroblast growth factor
CADD: Combined Annotation Depndent Depletion score
cDNA: Complementary DNA
CHX: Cycloheximide
DMEM: Dulbecco’s Modified Eagle Medium
FBS: Fetal Bovine Serum
GARS: Groningen Activity Restriction Scale
gDNA: Genomic DNA
gnomAD: Genome Aggregation Database
GWAS: Genome-wide association studies
hMSCs: Human Mesenchymal Stem Cells
IADL: Instrumental Activities of Daily Living
IBD: Identity-by-descent
LLI: Long-Lived Individuals
LLS: Leiden Longevity Study
LOD: Logarithm of the odds score
LOD_max_: Peak maximum LOD-score
MAF: Minor Allele Frequency
MEM α: Minimum Essential Medium Alpha
Merlin: Multipoint Engine for Rapid Likelihood INference
mESC: Mouse Embroyonic Stem Cells
MMSE: Mini-Mental State Examination
(NF)E: (Non-Finnish) European
Non-prot. Alt.: Non-protein-altering
NPL: Non-parametric linkage
PCR: Polymerase chain reaction
SAIGE: Scalable and Accurate Implementation of Generalized mixed model
Sanger seq.: Sanger Sequencing
SA-β-GAL: Senescence beta-galactosidase
SNP: Single-nucleotide Polymorphism
SNV: Single Nucleotide Variant
VEP: Variant Effect Predictor
WES: Whole-exome-sequencing
WGS: Whole-genome sequencing

